# Unmmaped RNA-Seq reads from inoculated sugarcane reveals long non-coding RNAs related to retrotransposons and suggest microbiome modulation

**DOI:** 10.1101/2025.09.28.679069

**Authors:** Joel Nunes Leite Junior, Helba Cirino, Priscila C. Antonello, Marcelo Marques Zerillo, Henrique M. Dias, Marie-Anne Van Sluys

**Affiliations:** GaTELab, Departamento de Botânica, Instituto de Biociências, Universidade de São Paulo - USP; Aggeu Magalhães Institute, Fiocruz Pernambuco; Departamento de Botânica e Ecologia, Instituto de Biociências, UFMT; Department of Agronomy, Horticulture and Plant Science, South Dakota State University, Brookings, SD, United States

**Keywords:** whole transcriptome, transposable elements, Acinetobacter, Fusarium

## Abstract

**Background:** Transcriptome studies have contributed to the understanding of protein coding and non-coding gene expression of several organisms. They also provide knowledge into host responses to pathogen infections. Not common yet is to detect transcripts from multiple interacting organisms present in a given sample. In addition, transcriptome studies with complex polyploid hybrid genomes and not completely sequenced, such as the one of sugarcane, remains a challenge. In such studies, a considerable set of reads may not align on any reference genome but still hold significant relevance to the study, allowing to gather information beyond the complex organism itself.

**Results:** A complete transcriptome analysis of sugarcane inoculated with a pathogen, with and without the addition of a beneficial bacteria, generated a subset of reads that did not map to their respective genome references. Here, we report a pipeline based on the assembly of a collection of unmapped reads that potentially interfere with the gene expression of the associated microbiome, as well as the identification of long non-coding RNAs related to transposable elements present in a host-pathogen interaction. In addition, the detection of transcripts of naturally occurring microorganisms in sugarcane allowed the identification of its microbiome at the species level. Further studies support changes in microorganism transcription levels according to the biotic stress the plant was conditioned.

**Conclusion:** A quick and practical pipeline is proposed to study unmapped reads to infer relevant information that would remain otherwise unnoticed. New sequences from hybrid sugarcane transcriptome, such as long non-coding genes related to known retrotransposons are described. Also, changes in microbiome gene expression provides insights to the microbiome alterations and bring knowledge to the sugarcane “pathobiome”.

## BACKGROUND

Studies based on high-throughput sequencing technology (HTS) allow inferring a great diversity of information based on the generated deep sequencing data (Reuter et al., 2015). Nowadays, it is possible to sequence a given sample from any organism and extract information from changes in expression levels of different genes after sequence alignment, similar to what is done in metagenome studies. Model organisms benefit from well-assembled and annotated reference genomes, which enable the identification of gene expression levels of protein-coding and non-coding genes, usually relying on sequencing messenger RNA by enriching the sample with a 3’ poly(A) tail captured by oligodT. However, this approach primarily restricts the sample to RNA polymerase II (RNA Pol II) transcribed genes and may leave relevant information unnoticed. In this context, the ribosome depletion of total RNA prior to transcriptome sequencing allows the disclosure of underrepresented sequences, such as non-coding RNA or non-polyadenylated transcripts, as opposed to other approaches such as mRNA-enrichment sequencing (mRNA-Seq), that is often and erroneously referred as RNA-seq (St Laurent et al., 2015).

Most transcripts in human cells belong to the class of non-protein coding genes (Bridges et al., 2021), a phenomenon that also occurs in other eukaryotes such as plants (Chekanova et al., 2007). Among the wide variety of non-coding RNAs there are the well-established class of RNA interference (RNAi), which has a double-stranded RNA as a precursor during its biogenesis and can be classified as microRNAs (miRNA), small interference RNAs (siRNA), and Piwi-interacting RNAs (piRNA) (Jarroux et al., 2017). Additionally, another class, long non-coding RNAs (lncRNAs), which are larger than 200 nucleotides, can have multiple functions, such as regulating developmental processes (Yu et al., 2019) in response to abiotic stress (Deng et al., 2019) The great diversity of lncRNAs allows their classification based on various criteria, such as transcript size, their relationship to protein-coding genes (PCG) or to other DNA elements, subcellular localization, or their function (St Laurent et al., 2015; Jarroux et al., 2017).

A whole-transcriptome approach also enables the identification of gene expression on host-associated microbiomes. Only recently, few reports have addressed the challenges and opportunities in this field (Liao et al., 2019; Westermann et al., 2017; Li et al., 2016) most using model organisms and focusing on the specificities of these associations. The present study describes a plant-microbe interaction, focusing on reads that did not map on the reference genomes of the three organisms used in this study.

High quality reads retrieved from the transcriptomes of *i*. non-inoculated sugarcane, *ii*. sugarcane inoculated with a pathogen, and *iii*. sugarcane coinoculated with pathogen and a beneficial bacteria, that failed to mapped to their reference genomes were assemble into contigs and used in further analyses.

This is a practical approach to identify unmapped RNA-Seq reads derived from the whole transcriptome of sugarcane inoculated with bacteria after sequence alignment to their respective genomes. It also enables the capture of diverse information about sequences from both the host and associated microorganisms interacting with the host. A total of 190 lncRNA were identified as belonging to the sugarcane genome and derived from transposable elements (TEs). Additionally, evidence of microbiome expression modulation between treatments was observed.

## RESULTS

### Unmapped reads assembly reveals informational transcripts in sugarcane

Paired-end high quality reads retrieved from three biological conditions, sugarcane mock inoculation (Sc), sugarcane inoculated with *X. albilineans* (ScXa), and with *X. albilineans* and *G. diazotrophicus* (ScXaGd), were aligned with their available genomic references. Paired-end reads with at least 80% coverage and 80% identity with the ribosomal gene clusters from sugarcane (nuclear, mitochondria, and chloroplast), *X. albilineans* (Miranda et al., 2023) and *G. diazotrophicus* (Bertalan et al., 2009) were filtered out. Reads were then compared to the genomes of *X. albilineans* and *G. diazotrophicus*, and removed if alignments had more than 80% coverage and 80% identity. The remaining reads were aligned to sugarcane genome references, and those showing more than 80% coverage and 95% identity were set aside for further analysis in a separate study. The remaining subset, which did not map to microorganism genomes and showed low similarity to sugarcane, was retained and used as input for the *de novo* assembly conducted in this study.

### *De novo* assembly of unmapped reads captures organismal diversity

The unmapped reads were compared among themselves and clustered for redundancy removal using CD-HIT, when overlapping 80% or more of their length and sharing at least 90% of nucleotide identity, yielding 2,790 contigs larger then 1,000 nucleotides. This subset was aligned using BLASTn against the Non-Redundant (NR) database with default parameters, and an initial taxonomic identification was assigned to each contig using the TaxonKit tool. Most of the contigs were related to plants, bacteria and fungi, while other were related to less frequent groups, such as Sugarcane bacilliform virus or Arthropoda (Figure 2). After identifying similar sequences using CD-HIT they were assigned to the different treatments assessed here. Interestingly, some transcripts of the bacterium *Acinetobacter soli* were assembled in the control sugarcane treatment and in the co-inoculation with *G. diazotrophicus*. However, most of the sequences shared across treatments were plant-related, with many corresponding to *Saccharum sp*., while others were associated with different grasses such as *Sorghum bicolor* and *Miscanthus floridulus*. To expand the view of the plant-pathogen interaction, subsequent analyses focused on contigs classified as belonging to Poaceae (1,540), Bacteria (417) and Fungi (562) contigs (Figure 2). Unmapped reads from the Poaceae group were analyzed to identify low-conservation genomic sequences, such as lncRNAs and TEs, while bacterial and fungal transcripts were used to detect non-inoculated microorganisms and their gene expression profiles.

### Detection of novel transcripts from Saccharum hybrid cultivar related to Poaceae

Out of a total of 1,540 Poaceae contigs, only 211 were automatically assigned to the *Saccharum officinarum* complex (Figure 2). Considering that, contrary to the virtual impossibility of maintaining an axenic environment for bacteria in host inoculation experiments, sugarcane was the only plant present in the assay, and a possible explanation for its underrepresentation in TaxonID is because most of sugarcane genome was not sequenced yet sequence (Gasmeur et al., 2018; Souza and Van Sluys et al., 2019) (Figure 1). Thus, to look for similarities of these possible genes with the genomes of the Saccharum group we analyzed the alignment profile of these transcripts in different sugarcane cultivars: *S. officinarum* LA-purple, *S. spontaneum* AP85-411, *S. spontaneum* NP-X, the diploid genome *S. rufipilum* (Wang et al., 2023), R570 (Healey et al., 2024) and the hybrid cultivar SP80-3280 (Souza & Van Sluys et al., 2019). The metric for assessing sequence similarity was calculated by multiplying the percentage coverage by the percentage identity of the first hit of each contig in the genomes. This result was then divided by the maximum possible value in a perfect alignment, which is 10,000 (obtained by multiplying 100% coverage by 100% identity). This creates a value ranging from 0 to 1, where 0 indicates no similarity and 1 indicates complete similarity. Figure 3a shows the heatmap distribution of this analysis, revealing highly similar contigs in all taxa. Some contigs exhibited low similarity values against the SP80-3280 cultivar but high values against *S*. officinarum LA Purple and R570. In contrast, a set of contigs showed low similarity across all genomes, despite being differentially expressed during inoculation conditions (Figure 3, b). Functional annotation of these contigs identified the NR-ARC domain as the most frequent (Figure 3, c). Among the 1,540 Poaceae contigs, 716 were predicted by functional annotation, while 824 remained unannotated. Further searchers for potential non-coding transcripts were conducted using non annotated contigs by EggNOG. Since EggNOG primarily annotates protein-coding genes and conserved functional domains, unannotated contigs are enriched for novel non-coding elements, including lncRNAs that lack protein-coding potential or conserved domain architectures.

**Figure 1:**
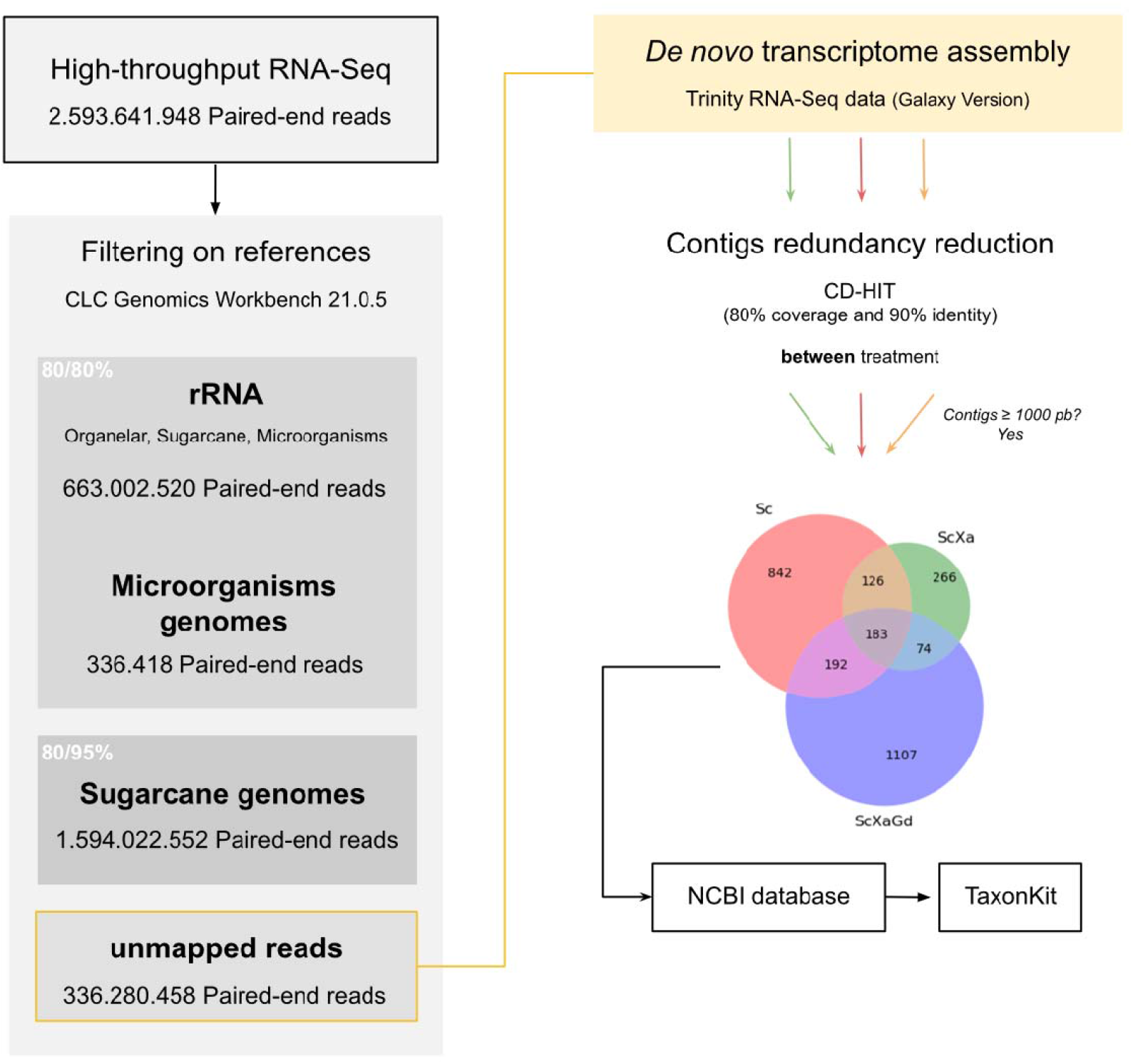
Pipeline for identifying unmapped sugarcane reads. After quality control, reads were mapped to the ribosomal gene clusters of known organism and genomic references, using alignment threshold of 80/80% coverage and identity for ribosomal sequences and bacterial genomes, and 80/95% for sugarcane genomic references. Paired-end (PE) reads that did not map in any of the reference sequences were used as input for the *de novo* assembly using Trinity. CD-HIT redundancy reduction, with 80/90% parameters resulted in 2,790 sequences longer than 1,000 nucleotides. These sequences were represented in a Venn diagram showing shared and unique transcripts among treatments. Furthermore, organisms were classified by searching the NCBI non-redundant database, and taxonomic classification was retrieved using TaxonKit.

**Figure 2:**
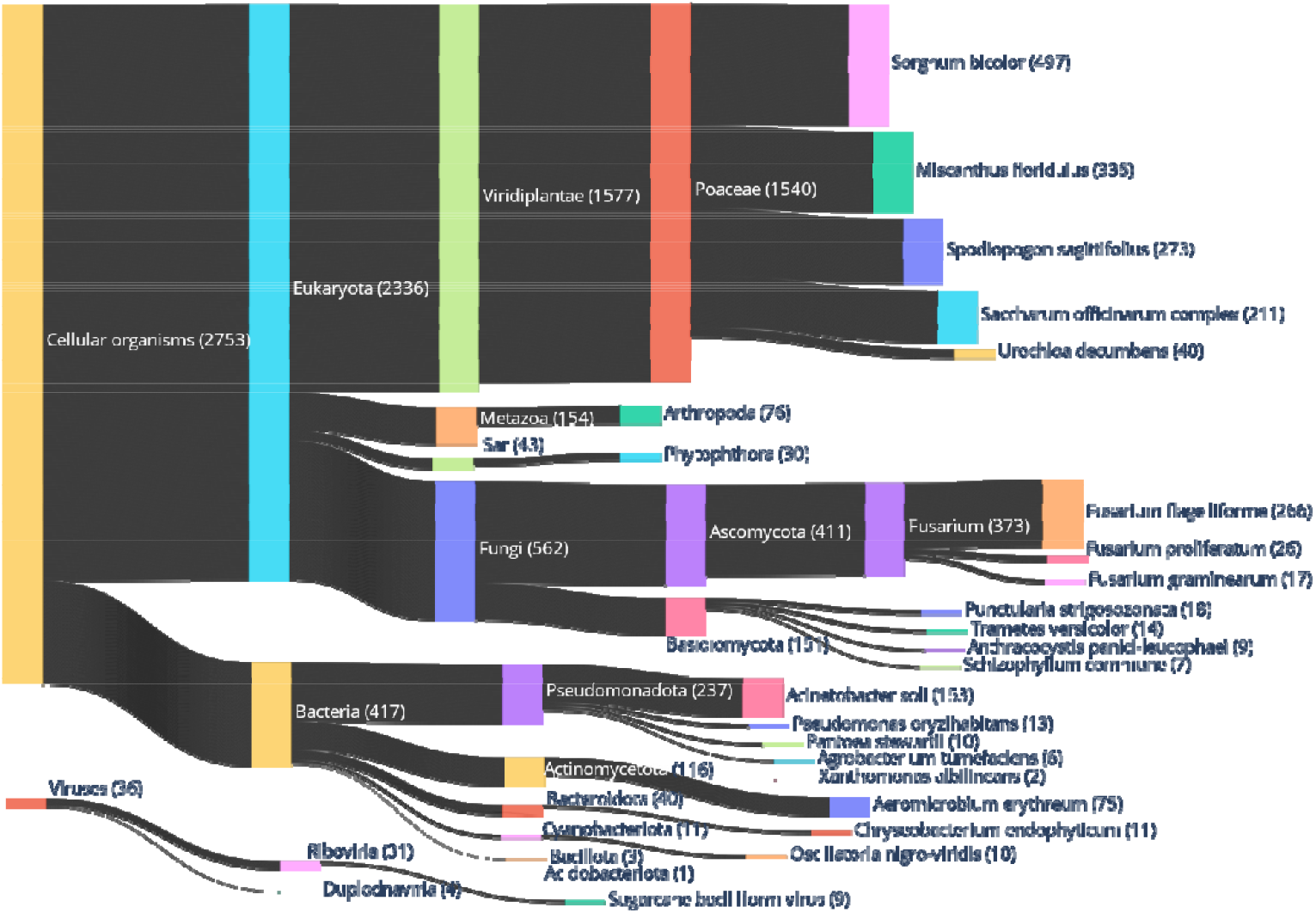
Taxonomic classification of the assembled contigs. The classification was performed by selecting the top hits of BLASTn and retrieving the corresponding TaxonID. Using this list of IDs, taxonomic classification was assigned using the TaxonKit tool. Of the 2,790 contigs, only one remained unidentified. Among the classified sequences, 1,540 sequences were classified as belonging to the Poaceae family, 417 to bacteria and 562 to fungi. Numbers in parentheses indicate the total contigs assigned to each organism. Contigs were identified at the species level for most organisms, with the greatest consensus observed for the bacterium *Acinetobacter soli*, which had 153 contigs assembled.

**Figure 3:**
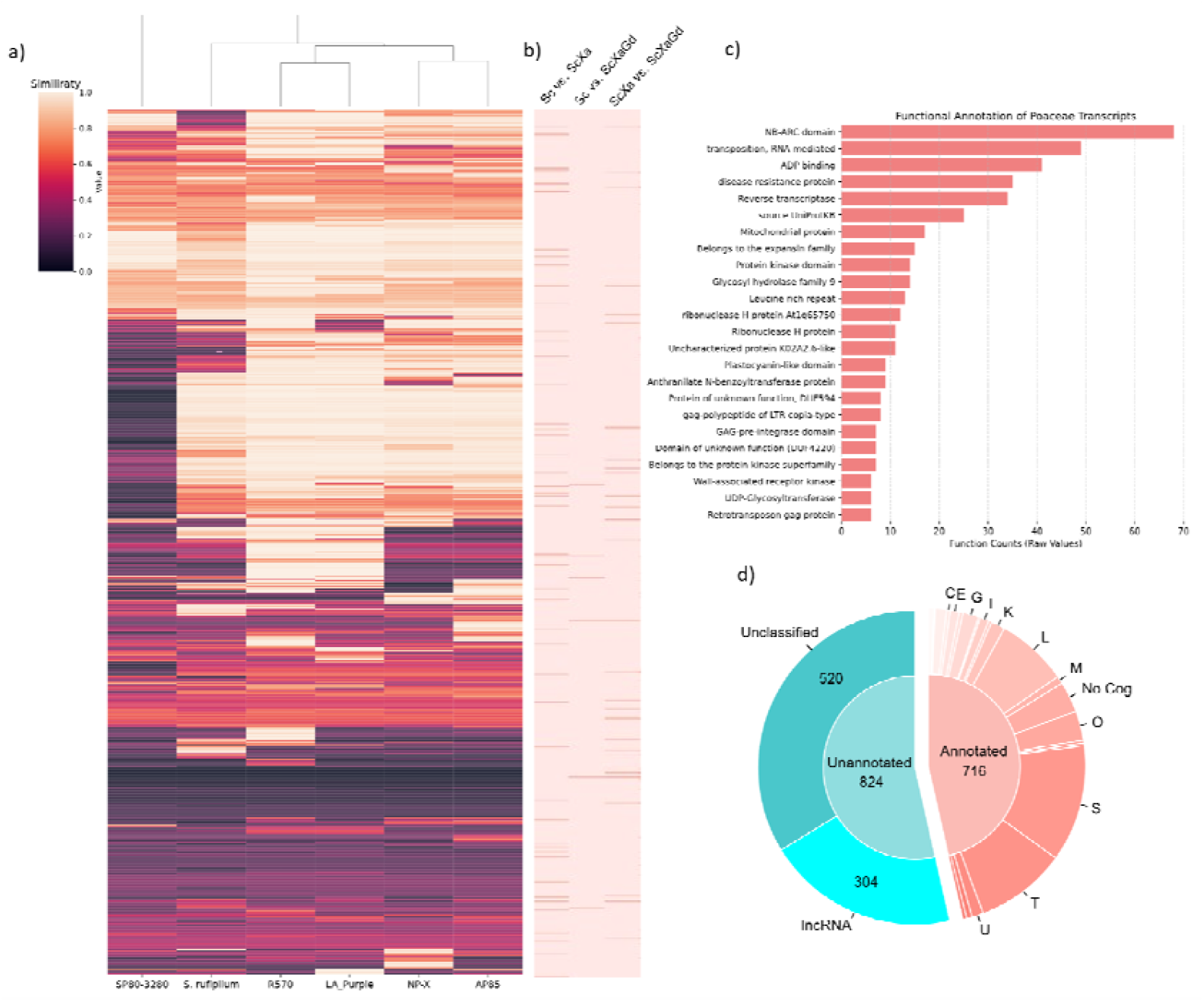
Features of Poaceae contigs. a) Heatmap showing megablast results of 1,540 contigs (rows) against five *Saccharum sp*. Cultivars (columns). Similarity index values were based on multiplying the percentage of coverage by identity, divided by the highest value obtained (100% x 100% = 10000) yielding an interval of 0 to 1 (dark to light). b) Poaceae genes quantified using Salmon quant and identified as differentially expressed across treatments using DESeq with p-adj < 0.05 and log2 FC ≥ 1. c) Functional annotation (Description field) and frequency of annotated genes of all treatment combined. d) Pie chart showing contig annotation results. EggNOG-annotated contigs (716) were classified into COG categories. Unannotated contigs (824) included putative lncRNAs (304) identified through subsequent analysis described in methodology.

Following an integrated lncRNA prediction pipeline either alignment-dependent or independent (Wang et al., 2023; Pooja et al., 2021; Jannesar et al., 2020), we prioritized the use of BASiNET that accepts new sequences and can tailor training to a non-model organism studied (Ito et al. 2018), as sugarcane. After training BASiNET with 2,600 coding sequences (CDS) and 2,600 non-coding sequences from *Sorghum bicolor*, achieving a true positive rate (TPR) of 0.985. Using this pipeline, we first classified the 824 Poaceae sequences that were not annotated by EggNOG. To avoid assuming a predicted lncRNA resulting from an overfitting, we performed BlastX searches of these 824 sequences against the Poaceae sequences from PDB, SwissProt, RefSeq and GenBank database. Only sequences with a maximum length of 100 amino acids, a size considered minimal for ORF annotation (Ruiz-Orera & Alba, 2019) were considered for the analysis and where many micropeptides derived from ncRNAs fit (Wu et al., 2022; Setrerrahmane et al., 2022). This process allowed us to predict 304 lncRNAs (Figura 3, d). Among these, 43 lncRNAs showed a differential expression during inoculations, with 14 showing differences exclusively in the Sc vs. ScXaGd comparison (Figure 3, b). We also identified 122 contigs annotated by EggNOG that were differentially expressed between treatments.

### Transposable elements related to unmapped reads

Although no differentially expressed genes (DEGs) was detected in this context, genes related to reverse transcriptase were predicted by functional annotation. Given that this study focuses on unmapped reads in the reference genome, we further explored whether these transcripts could be related to mobile genetic elements (retrotransposons and transposons). Thus, we used a custom-built sugarcane TE library (GaTE) from several previous studies (de Araújo et al. 2005, Domingues et al., 2012; Jesus et al., 2012; Saccaro-Jr et al., 2007) to identify whether these transcripts corresponded to previously described sugarcane TEs. After aligning Poaceae-related contigs against the multifasta TE database, we observed that of the 1540 Poaceae contigs, 305 contigs were related to TEs (threshold: ≥29 nucleotide alignment length), which were distributed across 13 TE families (Figure 4, a). Although some contigs aligned to multiple families or more than once within the same family, among the transposon class, the CACTA and hAT family presented the largest number of alignments. For LTR-retrotransposons classes, alignments over longer base pairs, mainly in the Gypsy superfamily, composed of the Del, Reina, TAT/Athila families. Significant coverage was also observed in the Copia retrotranposon superfamily, which includes Maximus/Sire, Ivana/Oryco, Angela/Tork, TAR/Tork. Among all TE families analyzed, DEL and Maximus/Sire stood out for presenting more alignments, despite Maximus/Sire aligning more times in different contigs (Figure 4, a).

**Figure 4:**
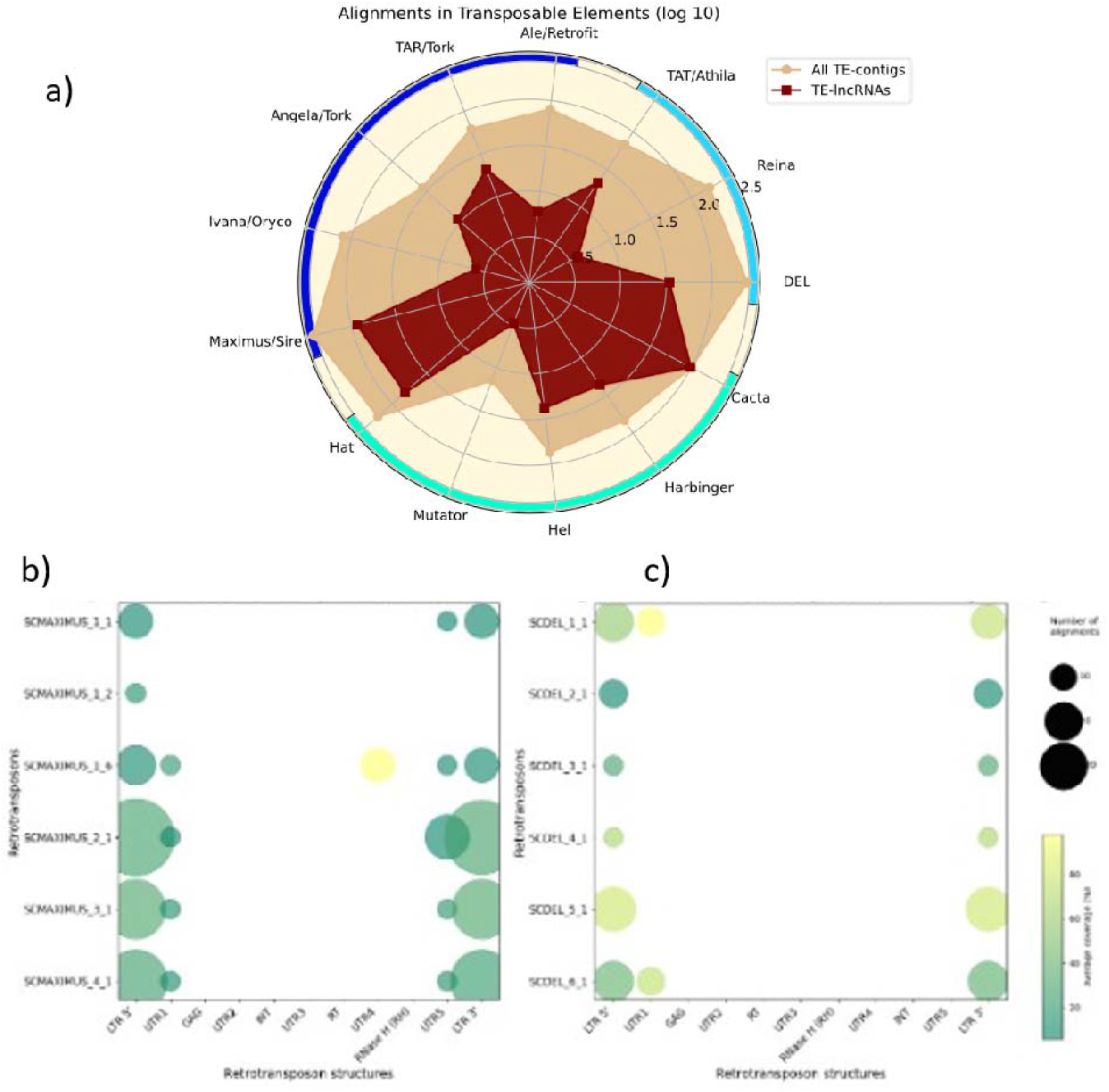
Transposon element-related sequence analysis. a) TE families aligned against the 1540 Poaceae sequences with their alignments and respective different contigs aligned. Colors in the outer circle indicate TE classes, being dark and light blue for LTR-retrotransposons, Copia and Gypsy, respectively, and green for transposon classes. b) Alignment of lncRNAs in Maximus and c) Del family lineages, respectively, in the respective TE structures.

We selected these two LTR-retrotranposon families, which had the highest number of alignments, to narrow down on TE classification regions where the contigs aligned. In addition, based on Domingues et al. (2012), we considered the lineages identified within the two families of LTR Retrotransposons in order to detect whether any lineage might show a different behavior from the others. Aligned transcripts were identified in some of the internal coding regions, such as the Gag and reverse transcriptase (RT) enzymes (Figure 2c). Knowing that LTR can act as promoters of non-coding RNAs (Cohen et al., 2009), and that we identified 304 lncRNAs from our 1540 Poaceae contigs (Figure 2, c), we sought to relate the sequences of long non-coding RNAs to these TEs. Thus, we recorded the information that of the 305 TE-related contigs, 58 were lncRNAs (lncRNA-TE) and when analyzing the alignment pattern of these lncRNAs, all of them showed a preference for the 5’ LTR, 3’ LTR, and untranslated regions (UTR) in DEL and Maximus family (Figure 4b, 4c). In addition, we used the library of small RNAs (sRNA) from TEs made available by Domingues et al. (2012) to investigate potential associationsbetween our lncRNAs and sRNAs. After performing a specialized blastn search for sequences shorter than 50 base pairs (-task blastn-short), we obtained 58 TE-lncRNAs out of all 304 previously identified that showed alignments against this sRNA library.

### Transcriptional modulation of the microbiome

Among the contigs with taxon identication, 417 were classified as bacterial, with 237 related to the phylum Pseudomonadota/Proteobacteria, although representatives from other phyla were also present (Figure 2). However, the frequency of assembled contigs identified in the NR database varied significantly by bacterial species, particularly for *Acinetobacter soli*, which accounted for 153 contigs, and *Aeromicrobium erythreum* with 75 (Figure 2), both representatives of the Pseudomonadota and Actinomycetota phyla, respectively.

Although redundancy was removed from the contigs across treatments, no contigs from *Acinetobacter soli* were assembled in the treatment with *X. albilineans* (ScXa). To better understand the distribution of RNASeq reads from detected bacteria among treatments, we aligned them against the reference genomes of the most abundant species from each genus, following a metatranscriptome approach (Chong et al., 2019; Law et al., 2022). Since some genomes contained multiple scaffolds and plasmids, we consolidated all sequences into a pseudochromosome to facilitate the analyses. To maximize the detection of bacterial reads in rRNA references and bacterial genomes (Figure 1), we used a threshold of 97% identity, previously considered a suitable benchmark for bacterial species classification (Schloss & Handelsman, 2005). Although this approach is no longer optimal for species-level identification (Edgar, 2018), it remains adequate for aligning genomic DNA reads within bacterial OTUs at the genus level (Dong et al., 2018), or for fungal community classification at 95% identity (Niu et al., 2015). However, in this study, we adopted a more stringent approach, aligning RNA-Seq reads in genomes with ≥ 99% identity and ≥ 95% coverage to increase accuracy while maintaining sensitivity for bacterial identification.

The results revealed patterns that distinguish the three groups Sc, ScXa and ScXaGd (Figure 5, a). It was observed that for some bacteria, there is a pattern of transcript abundance among the replicates of each treatment, but it varies depending on the treatment. For example, transcripts derived from *Acinetobacter soli* were abundant in the control (Sc), followed by sugarcane co-inoculated (ScXaGd), while were underrepresented in sugarcane with the presence of the pathogen (ScXa). In contrast, *Pantoea stewartii* related transcripts were poorly detected in the control, and consistently abundant in all replicated of ScXa (Figure 5, a). On the other hand, transcripts aligned to *Pseudomonas oryzihabitans* showed statistically significant (pvalue < 0.05) higher total counts in the ScXaGd treatment, while there is a significant decrease in the ScXa treatment. Although *A. soli* did not exhibit statistically significant differential expression between treatments, it had the highest number of assembled contigs in this set.

**Figure 5:**
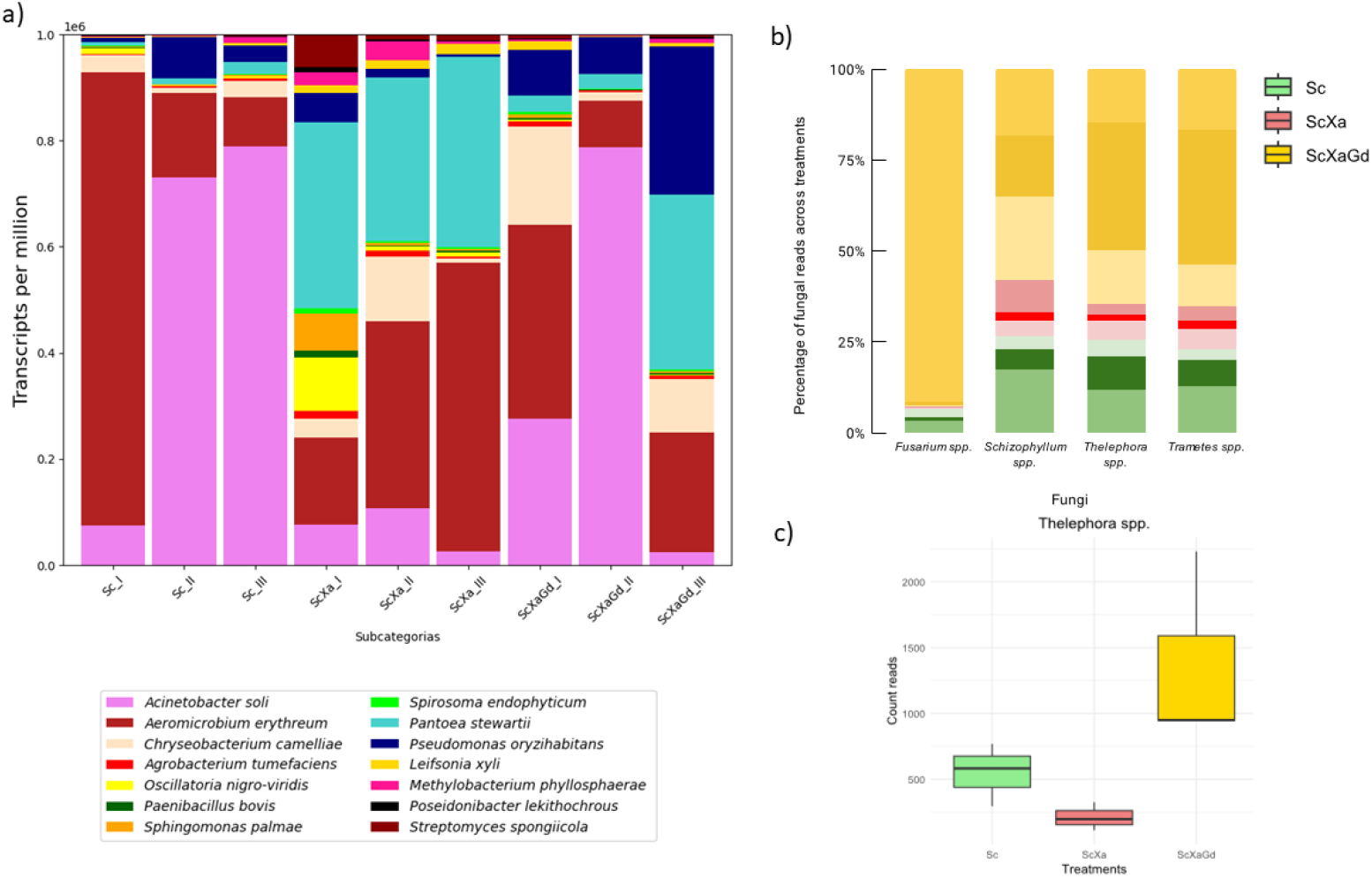
Modulation of the microbiome at the transcriptional level. a) The coverage of RNASeq reads in bacterial genomes obtained from GenBank was calculated (Transcripts Per Million) and the distribution of the 14 bacteria that got more than two contigs assembled was plotted. Alignment parameters: 90% coverage and 99% identity. b) Following the read alignment in selected fungi genomes, a straightforward normalization was conducted using the percentage of read counts in each library relative to the total counts across all treatments. Shades of the same color represent the different replicates for each treatment. Alignment parameters: 90/95% for coverage and identity, respectively. c) Total counts reads for *Thelephora spp*. with significant difference between treatments performing Dunn’s test with Bonferroni correction between ScXa and ScXaGd with p-value less than 0.05.

To further investigate, we analyzed which genes these contigs correspond to. From 153 *A. soli* contigs (Figure 2), none were assembled in ScXa treatment, but we identified two candidate genes: glutamine synthetase (*glnA*), only expressed in the Sc control and Lipopolysaccharide-assembly, LptC-related (*lptC*), which was only expressed in the ScXaGd group. Additionally, DEG analysis revelead that iron-sulfur (Fe-S) clusters (*iscS*) were significantly more expressed in Sc compared to ScXaGd. As well as observing a number of contigs related to bacteria, we identified 562 fungal-related contigs, with 373 associated with *Fusarium spp*. (Figure 2). Notably, most fungal contigs were assembled with reads from the ScXaGd treatment. To further examine fungal transcript distribution across treatments, we applied the same approach used for bacterial profile analysis (Figure 5, a). Due to the inconsistencies in species specificity within the same genus, we relaxed the RNASeq alignment parameters to 90/95% to enrich the genera of these genomes and perform a more comprehensive analysis. After normalizing the percentage of read counts in each library relative to total counts of all treatments, a concentration of fungi reads was observed in the ScXaGd treatment for all genus fungi analysed (Figure 5, b), being statistically significant (pvalue < 0.05) for *Thelephora spp*. in relation to ScXa (Figura 5, c).

## Discussion

### Unmapped reads as a source of novel insights into the sugarcane transcriptome

The present work describes a strategy to analyze unmapped reads from whole transcriptome derived from host-pathogen interaction. As described, reads were sorted according to the genomes of known interacting organisms. Achieving comprehensive coverage of both the eukaryotic host and prokaryotic pathogen enables us to examine metabolism from a pathogenic perspective and confirms the efficacy of the methodology described previously (Zerillo et al., submitted).

From the unmapped reads, we obtained a diverse set of assembled contigs, including potential transcripts from sugarcane hybrid cultivar used. Similar to maize (*Zea mays*), sugarcane (*Sacharum spp*.) belongs to Poaceae family. However, while maize is a diploid with a 2.4 Gb genome (Haberer et al., 2005), sugarcane has a much larger (∼10 Gb) and polyploid genome. Parental species such as *Saccharum officinarum* and *Saccharum spontaneum* have genomes reaching about 8.55 Gb and 12.64 Gb, respectively (Zhang et al., 2012), while hybrids cultivar can reach even higher ploidy levels. Several reference genomes exist for sugarcane cultivars, including the assembled genome of R570 (Healey et al., 2024), an allele-resolved genome of the hybrid commercial sugarcane cultivar SP80-3280 (Souza, Van Sluys et al., 2019), and tetraploid *S. spontaneum* genomes (Zhang et al., 2018), along with a diploid reference (Wang et al., 2022), but none represent the completion of commercial sugarcane genome, which may be represented approximately by 40%, when all references are combined.

Since all references used in this study are genomic, while our data is transcriptomic, it is expected that complete matches would be absent due introns removal, RNA polymerase errors (Cheung et al., 2020), or alternative splicing events (Lam et al., 2022), which may occur in response to abiotic stresses (Yang et al., 2022) or microorganism interactions (Rigo et al., 2019). Greater similarity was observed with the *S. officinarum* cultivar LA Purple compared to other *S. spontaneum* derived varieties. This is expected, as R570 cultivar has only ∼10% *S. spontaneum* content, with the remainder originating from *S. officinarum* or recombinant chromosomes (Cuadrado et al., 2004). The low similarity of some contigs across all sugarcane cultivars suggests that these sequences may be underrepresented in current genomic references. Given that SP78-4467 is a susceptible cultivar, it is conceivable that it shares less similarity with *S. spontaneum*, whichis know to contribuite resistance genes in sugarcane breeding programs (Souza and Van Sluys et al., 2019).

### Functional annotation of host-pathogen interaction genes and lncRNAs

Functional annotation of the putative genes indentified NR-ARC Domain resistance genes (Pan et al., 2022) and other defense-related gene sequences such as Expansin family related (Backiyarani et al., 2022). Additionally, genes involved in pathogen-plant interaction were differentially expressed such as lipoxygenase (Camejo et al., 2019), GDSL esterase/lipase (Vincent at al., 2022) and low sulfur response genes (Mukhtar, et al., 2011). Of the 1,540 Poaceae contigs, 824 were not annotated by EggNOG, of which 304 were classified as lncRNAs, an expected finding since conservation is not a typical characteristic of lncRNAs (Huang et al., 2023), particularly in a dataset derived from a subset of unmapped reads. The search for lncRNAs was crucial for understanding some of the differentially expressed transcripts, as 42 of them were identified as potential lncRNAs that were not detected through functional annotation or BLASTx analysis.

### Transposable elements and their relationship with lncRNAs

Functional annotation also revealed reverse transcriptase activity, suggesting active retrotransposons (Moran & Wilson, 2022). In the human genome, nearly of the half of the sequences is composed of TE insertions, though most are no longer active (Chénais, 2022), while in plants, TEs are even more abundant, constituting up to 80% of some genomes, as is the case in maize (Zhang & Qi, 2019). To investigate this, instead of using Repbase Update (Bao et al., 2015), we used a curated sugarcane TE database previously identified (Jesus et al., 2012; Setta et al., 2012; Domingues et al., 2012; Rossi et al., 2004). Many contigs contained only partial retrotransposon insertions, corroborating the idea that TEs contribute to the generation of mutations and genomic rearrangements (Bourque et al., 2018). The relationship between TEs with lncRNA is already established in plants (Yadav et al., 2023; Lv et al., 2019; Wang et al., 2017), which TEs are known to give rise to non-coding RNAs (Bourque et al., 2018), including retrotransposons have been described (Ganesh & Svoboda, 2016). For instance, the LINE retroelement generates a lncRNA in human (Cartault et al., 2012) and octopus brains (Petrosino et al., 2022). In plants, lncRNAs can also be derived from retrotransposons (Wang et al., 2017), for example, being derived from the Gypsy family (Li et al., 2022) or derived specifically from the LTR region (Kirov et al., 2020) and include small conserved fragments that could serve as an active site for production or target of sRNAs as some study suggest (Yan et al, 2020; Zheng et al., 2022; Huo et al., 2022).

### Microbiome composition and host influence on microbial communities

In addition to Poaceae 1,540 sequences, we identified 417 and 562 contigs referring to bacteria and fungi, respectively, allowing for functional-level exploration of the sugarcane microbiome. Interestingly, we observed that among the most abundant microbial taxa belonged to the phylum Pseudomonata/Proteobacteria (bacteria) and Ascomycota (fungi), consistent with findings by Trivedi et al. (2021), who described similar profiles of bacterial and fungal communities in the leaf endospheres. Other studies on sugarcane microbiome, also have reported *Pseudomonas sp*., *Acinetobacter sp*., *Streptomyces sp*., *Sphingomonas sp*. and *Pantoea sp*. in sugarcane roots (Dong et al., 2018), while *Chryseobacterium sp*. and *Agrobacterium sp*. were also found in bulk soil of sugarcane roots (Yeoh et al., 2015). Interestingly, Ishida et al., (2022) identified similar genera in two additional cultivars and documented their relative abundance in different plant compartments and organs. These genera included *Methylobacterium sp*., *Pantoea sp*., *Pseudomonas sp*., *Acinetobacter sp*., and *Leifsonia sp*., suggesting that microbial communities associated with sugarcane may be conserved across different genotypes, regardless of susceptibility or resistance.

The influence of the host genome on the microbiome has been widely discussed (Tabrett & Horton, 2020) in both animals (Wang et al., 2016) and plants (Chen et al., 2020). In rice, Zhang et al. (2022) identified that the microbiome is shaped by microhabitats and contains several of the same bacterial genera detected in our study. However, we provide evidence that microbiome modulation is driven by pathogen inoculation, supporting the hypothesis that disease phenotypes arise not only from pathogen itself but from the pathobiome, a term that defines the entire disease-related microbial community (Defazio et al., 2014; Mannaa & Seo, 2021).

### Microbial modulation and functional insights

We further reveal the use of transcriptome as a tool for identify an *A. soli* contig referring to the glutamine synthetase gene, one of the enzymes necessary for nitrogen assimilation (Liu et al., 2022). This gene was expressed only in Sc control but absent in ScXa and ScXaGd, possibly due to nitrogen fixation caused by *G. diazothrophicus*, which may mitigate or inhibit the need for N fixation by *A. soli*. On the other hand, the lipopolysaccharide-assembly gene (*lptC*), associated with interbacterial competition (Trotta et al., 2023) was exclusively expressed in the treatment ScXaGd. *G. diazotrophicus* produces gluconacin, which has antagonistic activity to phytopathogens and some beneficial bacteria (Oliveira et al., 2018). This explains the reduction in transcripts from potentially pathogenic bacteria, such as *Pantoea stewartii*, but does not apply to beneficial bacterium *A. soli*, perhaps because *A. soli* can react well to competition, although studies on the interaction of these bacteria are needed.

The study of the microbiome also allowed us to explore bacterial-fungi interactions. For example, *Schizophyllum sp*., known to colonize various plants, inducing white rot disease, and facilitating lignin degradation with biotechnological applications (Tovar-Herrera et al., 2018; Kumar et al., 2022), has also been reported to exhibit increased growth in the presence of *Pantoea agglomerans* (Krause et al., 2020). Conversely, *Trametes sp*. contribute to white rot disease through the production of laccases that oxidize lignin-related compounds (Abbas et al., 2023), a mechanism also carried out by *Pantoea ananatis* (Zhang et al., 2018). Moreover, *Pantoea sp*. have been associated with an inhibitory activity against Fusarium spp. (Nakkeeran et al., 2021; Pandolfi et al., 2010), though *G. diazotrophicus* did not suppress Fusarium spp. presence in our dataset, as previously observed (Logeshwarn et al., 2011).

### Final considerations

Our approach allowed us to demonstrate the diversity of interactions that may be occurring in the plant during biotic stress at lncRNAs and microbiome level as a result of the imbalance of a taxonomic group, allowing us to further study the relationships between these microorganisms to understand their biological role in the face of each situation. In addition to the restrictive similarity parameters, we excluded contigs considered redundant from the whole analysis. Nevertheless, it was possible to infer information about why sugarcane reads do not align in our references, to detect long non-coding RNAs, differentially expressed contigs between treatments and related to transposon elements, and to detect transcripts of possible bacterial species that were having their transcriptional profile modulated as a function of biotic stress. In this context, the study of unmapped reads is highly relevant to assimilate and cover all the information that a total transcriptome can offer.

Once considered non-informative reads, our study reveals the potential of an unmapped RNA-seq dataset, especially when targeting the complex sugarcane genome. We demonstrate the ability to capture information that would otherwise go unnoticed by developing a pipeline with different identity and coverage parameters from those previously described. Thus, *de novo* assembly of unmapped reads allowed us to identify the similarity of these contig transcripts with part of the *Saccharum* group and to start extracting information about similarities with this susceptible cultivar. Three hundred four contigs were classified as long non-coding RNA, being differentially expressed between treatments. Interestingly, some of the *Saccharum* contigs were fully or partially aligned to previously described retrotransposons and transposons, indicating that they may be derived from new genomic regions not present in other cane cultivars. The assembled bacteria contigs disclose the presence of *A. soli* in this dataset among the bacteria. Interestingly, a change in the microbiome gene expression is observed revealing a transcriptional modulation of metabolic features. Also, changes in the microbiome prevalence are revealed, for instance, *P. stewartii* being more abundant under the pathogen-inoculated condition as well detection of fungal transcripts in coinoculation with *G. diazotrophicus*. Taken together, our data strengthen the importance of studying the unmapped reads to understand more deeply plant-microbe interactions.

## Material and Methods

### Biological material

The susceptible sugarcane cultivar SP78-4467 was used in a pilot experiment of interaction with microorganisms and conditioned to three treatments. Besides the control (Sc), without inoculation of bacteria, there is the condition of the presence of the pathogenic bacterium *Xanthomonas albilineans* Xa11 (ScXa) (Miranda et al., 2023; Tardiani et al., 2014) and in the presence of *X. albilineans* and *Gluconacetobacter diazotrophicus* PAL5 (ScXaGd) (Bertalan et al., 2009). Each treatment had three biological replicates, although the third biological replicate (iii) was distributed over two sequencing runs, with the total transcriptome extracted and rRNA depleted for subsequent sequencing.

### Sequencing and filtering

Performed with Illumina high-throughput sequencing technology for all biological replicates. However, the third replicate from each treatment was distributed over two sequencing runs, generating four lanes. With the sequenced libraries, different references were used, filtering each sequence and retaining the aligned reads by the RNA-Seq Analysis tool of CLC Genomics Workbench 21.0.5. For rRNA sequences and bacterial genomes, the parameters of 80% coverage and identity were used for the alignment of the reads in the sequence, while for Saccharum sp references it was 80% coverage and 95% identity. After all filtering, each replicate generated an amount of unmapped reads, which make up this paper.

Paired-end and high quality reads were first aligned with the available genome references of the three organisms. First mapping against sugarcane sequences, such as: plastidial rRNA operons of *Saccharum* sp. SP80-3280 (NC_005878.2) and Q155 (NC_029221.1); mitochondrial rRNA operons of *S. officinarum* Khon Kaen 3 (LC107874.1) and *Saccharum* sp. (MG969495.1); chromosomal rRNA operon of *Saccharum* sp. R570 (KF184927.1); and genomic fragments comprised of 4.26 Gb from SP80-3280 cultivar (QPEU01000000); and 382 Mb from 570 cultivar. Secondly, the non-mapped reads were then compared to *X. albilineans* strain Xa11 rRNA operons (GCA_030864115.1), Xa11 genome (GCA_030864115.1), and *Gluconacetobacter diazotrophicus* strain PAl 5 (AM889285, AM889286 and AM889287).

### *De novo* assembly and redundancy reduction

Trinity de novo assembly (v2.14) of RNA-Seq data was used to assemble the contigs with the unmapped reads, considering the default. We used the CD-HIT Suite: Biological Sequence Clustering and Comparison for exclusion of redundant sequences with the cd-hit-est package in the default. The files with contigs for each treatment were turn into just one file for redundancy reduction in the trancriptome. Subsequently, we pooled all the clusters with sequences larger than 1000nt from the three treatments to identify similar sequences between the treatments and added the coverage parameter (aL and aS) of 80%, while the identity was kept at 90% of the default. This was our input to the Venn’s Diagram.

### Differential expression

The values of total counts reads and transcripts per million (TPM) were obtained by the RNA-Seq Analysis tool of CLC Genomics Workbench 21.0.5. The alignment parameters of the reads were 90% coverage and 95% for the aligned reads in Saccharum contigs and 95% coverage and 99% identity for the aligned reads in bacterial genomes with respectively number accession: *Acinetobacter soli* (ASM1813912v1), *Aeromicrobium erythreum* (ASM150940v1), *Chryseobacterium camelliae* (ASM277059v1), *Agrobacterium tumefaciens* (ASM366790v1), *Oscillatoria nigro-viridis* (ASM31747v1), *Paenibacillus bovis* (ASM142101v2), *Sphingomonas palmae* (GCF_900109565.1), *Spirosoma endophyticum* (GCF_900112365.1), *Pantoea stewartii* (ASM1104447v1), *Pseudomonas oryzihabitans* (ASM766563v1), *Leifsonia xyli subsp. Xyli* (ASM47077v1), *Methylobacterium phyllosphaerae* (ASM193617v1), *Poseidonibacter lekithochrous* (ASM1328383v1) and *Streptomyces spongiicola* (ASM312236v1). In the analysis of fungi, we used 90% coverage and 95% identity with the genomes of *Fusarium oxysporum* (GCF_000271745.1), *Schizophyllum commune* (GCF_000143185.2), *Thelephora terrestris* (GCA_015956445.1) and *Trametes versicolor* (GCF_000271585.1). For the counts aligned in genomes, we used the Levene, Kruskall-Wallis statistical tests and Dunn’s post-hoc comparisons to analyze data, evaluating the homogeneity of variances, comparing medians between groups and identifying specific differences. For counts in contigs assembled, we quantified with Salmon (-l ISR) and used DeSeq2 p-adjust less than 0.05 and log2FC more than | 1 |.

### Identification of coding and non-coding RNAs

We based ourselves on three ways. First, we used sequences not annotated by EggNOG. Then, we use an alignment-free classifier based on complex networks and extracting features from network topologies, the Biological Sequences NETwork (BASiNET) (Ito et al., 2018) which was trained with 2600 coding sequences (cds) from *Sorghum bicolor* obtained from PHYTOZOME. The 2600 lncRNAs of *Sorghum bicolor* were obtained from CANTATA DB. This set was used to classify all 1540 Poaceae sequences. After 10 Fold Cross Validation, our training using BASiNET obtained a TP Rate of 0.985, FP 0.015 and F-Measure of 0.985. Furthermore, the intersection of these two paths was BLASTx against Poaceae sequences obtained from NCBI databases (PDB, SwissProt, RefSeq, GenBank) to avoid assuming false positives and tolerate a maximum of ORFs of up to 100 amino acids.

## Data availability

The raw transcriptome sequencing data (RNA-seq) is available under the NCBI-SRA submission code <u>SUB14793753</u>.

## Acknowledgments

This work was supported by grants to MAVS from FAPESP (2016/17545-8) and CNPq 310779/2017-0. JNLJ is recipient of a CAPES or CNPQ and FAPESP Scholarship (2023/08813-2); MZ is recipient of a FAPESP fellowship (2018/23646-7); HC was recipient of a FAPESP fellowship (2022/10296-3); PA was recipient of a FAPESP fellowship (2022/09736-9) and HMD (FAPESP 2019/08239-9).

## Author contributions

JNJR designed and performed the bioinformatic data analysis and wrote the manuscript; MZ contributed to mapping reads on references and help to contigs assemblies; HC performed the statistical analysis of the differentially expressed contigs and assisted in the training stage; PA helped with statistics methods; HMD discussed result; and MAVS designed and coordinated the study, analyzed results and wrote the manuscript.

